# CYCLODEXTRIN DERIVATIVE ENHANCES THE OPHTHALMIC DELIVERY OF POORLY SOLUBLE AZITHROMYCIN

**DOI:** 10.1101/2020.06.16.154351

**Authors:** Anil Thakur, Sourabh Jain, Anjali Pant, Akanksha Sharma, Rajiv Kumar, Gajanand Sharma, Neha Singla, Ashish Suttee, Santosh Kumar, Ravi Pratap Barnwal, O.P. Katare, Gurpal Singh

## Abstract

Azithromycin (AZM), a macrolide antibiotic used for the treatment of Chlamydial conjunctivitis, is less effective for the treatment of this disease due to its poor bioavailability (38%). Various alternatives have been developed for improving the physico-chemical properties (i.e., solubility) of the AZM without much success. To overcome the problems associated with AZM, an inclusion complex employing a modified cyclodextrin i.e., sulfobutylether-β-cyclodextrin (SBE-β-CD) was prepared and characterized by phase solubility studies, pXRD and FTIR techniques. The results portrayed the formation of the inclusion complex of AZM with sulfobutylether β-cyclodextrin (SBE-β-CD) in 1:2 molar stoichiometric ratios. This inclusion complex was later incorporated into a polymer matrix to prepare an *in situ* gel. Various combinations of carbopol 934P and hydroxypropyl methylcellulose (HPMC K4M) polymers were used and evaluated by rheological and *in vitro* drug release studies. The optimized formulation (F4), containing carbopol 934P 0.2% (w/v) and HPMC K4M 0.2% (w/v), was evaluated for clarity, pH, gelling capacity, drug content, rheological properties, *in vitro* drug release pattern, ocular irritation test and antimicrobial efficacy. Finally, owing to the improved antimicrobial efficacy and increased residence time, AZM:SBE-β-CD *in situ* gel was found to be a promising formulation for the efficient treatment of bacterial ocular disease.

## 1. Introduction

The eye being one of the most important parts of the body has been the subject of intense research since a very long time [1, 2]. Many forms of eye-related infections are caused by pathogenic bacteria, but few to no feasible and viable treatments exist for this. Investigation of various ophthalmic dosage forms, like solutions, ointments, gels and polymeric inserts has been carried out in order to prolong the ocular residence time of medications intended for topical application to parts of the eye[3]. These dosage forms have been successful in increasing the corneal contact time to varying degrees. One of the major challenges faced in the delivery of ophthalmic drugs is the attainment and retention of optimal concentration of drugs at the site of action within the eye. within the eye. Further, ophthalmic preparations have less patient compliance because of blurred vision (e.g. ointments and inserts)[4].

The conventional systems such as eye drops suffer major drawback of having very poor bioavailability due to the rapid washout during lachrymation in eyes[5].The anti-microbial efficacy of moxifloxacin by forming an *in situ* gel and administering via ophthalmic route has reported the antimicrobial effect of moxifloxacin by forming an *in situ* gel upon ophthalmic administration. This resulted in gelation after instillation into the eye due to physico-chemical changes *(i.e.,* pH), thereby increasing the pre-corneal residence time and overall bioavailability of drug into the eye [6].Hence, it was observed that the sol-to-gel phase transition of the *in situ* gel on the eye surface would be dependent upon use of different polymeric composition and their structural behavior at varied pH ranges. In addition the sol-gel formation was proven to be effective by usage of thermo-sensitive, ion-activated, electric-sensitive, magnetic field-sensitive, ultrasonic-sensitive and chemical material-sensitive approaches. Subsequently, many researchers have reported various systems such as i) pH-trigger (e.g. cellulose acetate hydrogen phthalate latex), ii) temperature dependent (e.g. pluronics and tetronics), and iii) ion activation (i.e., gelrite)[7] for gel formation. In fact, another group had prepared *in situ* gel for ophthalmic preparation containing ciprofloxacin hydrochloride that was successfully formulated as *in situ* gel-forming eye drops with the help of sodium alginate and hydroxypropyl methylcellulose (HPMC)[8]. These results demonstrated that the alginate and HPMC mixture could be used as an *in situ* gelling carrier to improve ocular bioavailability and patient compliance. Moreover, various antimicrobial agents have poor bioavailability leading to inadequacy in the therapeutic regime of the eyes. In this study we have focused on Azithromycin (AZM), semi-synthetic, acid-stable macrolide-antibiotic. It has a long half-life (t_½_= 68 hours) and good tissue penetrability therefore, representing a profound alternative for the treatment of odontogenic infections. In addition AZM has a wide antimicrobial spectrum of action towards gram-negative bacilli, including anaerobic bacteria. It is effective against periodontal pathogens like *Aggregatibacter actinomycetemcomitans* (previously *Actinobacillus actinomycetemcomitans)* and *Porphyromonas gingivalis* and various studies support its use for the treatment of periodontal infections [9]. Sulfobutylether β-cyclodextrin (SBE-β-CD), a β-cyclodextrin derivative has been used before as a formulating agent to increase the solubility of sparingly soluble compounds. Hence, the present work attempts and focuses on the preparation, characterization & evaluation of AZM:SBE-β-CD complex and their further incorporation in gel for improving ophthalmic delivery. It is based on the concept of pH-triggered *in situ* gelation for ophthalmic delivery of AZM. The polymer polyacrylic acid (carbopol 934P) was used as the gelling agent since it exhibits a pH dependent sol-gel phase transition. Further, the AZM:SBE-β-CD inclusion based *in situ* gel may prove to be a feasible and viable substitute to traditional eye drops by imparting prolonged pre-corneal residence time and sustained released characteristics to AZM.

## 2. Materials and Methods

### 2.1 Materials

AZM was procured ex gratis from Sun Pharma, Poanta Sahib,India. Carbopol 934P (Loba Chemicals, Mumbai), HPMC K4M (Colorcon), SBE-β-CD (Cydex Pharmaceuticals,), Edetate disodium (Loba Chemicals, Mumbai), Benzalkonium chloride (Ases chemicals works, Jodhpur), Citric acid (Ases chemicals works, Jodhpur), Disodium hydrogen phosphate (Loba Chemicals, Mumbai), Whatman’s filter paper-41 (Whatmann Int. Ltd, England), Purified water (In house Laboratory) were used as it is for the experiments. Albino rabbits were obtained from the central animal house, Lachoo Memorial College of Science and Technology (Autonomous), Jodhpur, Rajasthan for the testing.

### 2.2 Methods

#### 2.2.1 Preparation of AZM: SBE-β-CD inclusion complex

A physical mixture of AZM with SBE-β-CD was prepared in a 1:2 molar ratio using a mortar to obtain a homogeneous powder blend. Inclusion complex of AZM:SBE-β-CD was prepared by kneading method[10]; appropriate amounts of AZM:SBE-β-CD in 1:2 M ratio were kneaded to a paste-like consistency with small amount of purified water. After grinding the paste it was evaporated under reduced pressure and dried in a vacuum oven to obtain dried powder of AZM:SBE-β-CD inclusion complex.

#### 2.2.2 Physicochemical Characterization

##### 2.2.2.1 Powder X-ray diffraction study (PXRD)

X-ray diffraction is a technique used to determine the atomic and molecular structure of a crystal wherein a beam of incident X-rays is caused to diffract into various distinct directions by the crystalline atoms. The powder X-ray diffraction (PXRD) patterns were determined for AZM, SBE-β-CD and AZM:SBE-β-CD physical mixture and their inclusion complex with diffraction angles (2θ) i.e., 2°-40° for the characterization of their structural properties[11].

##### 2.2.2.2 FTIR spectroscopy

The IR absorption spectrum of AZM, SBE-β-CD was recorded using dispersive powder technique scanned in the range of 4000-1000 cm^-1^. FTIR analyzes chemical bonds within a molecule; the spectrum generates a sample profile in the form of a distinguishable molecular fingerprint that can be used to screen and scan samples for many different components. The samples of AZM, SBE-β-CD, AZM:SBE-β-CD inclusion complex and AZM with SBE-β-CD physical mixture were pressed with potassium bromide (KBr) to form pellets. FTIR were recorded at frequencies from 4000-1000 cm^-1^ with resolution of 4 cm^-1^.

##### 2.2.2.3 Analytical Method development for Azithromycin using UV-visible spectroscopy

For determination of absorption maxima (λ_max_), 10 mg drug was dissolved in 100 ml phosphate buffer (pH 7.4) to prepare the stock solution (100 μg/ml). Subsequently, different concentrations (30, 40, 50, 60, 70, 80, 90 and 100 μg/ml) of drug were prepared by transferring 3,4,5,6,7,8,9 and 10 ml of the stock solution in 10 ml volumetric flask and volume made up with phosphate buffer (pH 7.4). The λm_ax_ was found to be 232 nm. Further on, the absorbance values of these solutions were taken at λ_max_=232 nm using UV-visible spectrophotometer (Shimadzu UV-visible spectrophotometer 1400).

##### 2.2.2.4 Solubility determination

The AZM solubility was determined by shake flask method. An excess quantity of AZM was added in 10 ml of phosphate buffer (pH 7.4) (supersaturated) and stirred at room temperature for 72 hrs until the equilibrium was attained. The solution was then passed through 0.45 μm membrane filter and the amount of the drug dissolved was analyzed by UV-visible spectrophotometer as before [12].

##### 2.2.2.5 Phase solubility studies

Standard solution of SBE-β-CD was prepared by dissolving 2 g in 10 ml of water (185 mM). Simultaneously, various dilutions of concentrations 46 mM, 23 mM, 11.6 mM and 0 mM were prepared. Further, AZM was added in all the diluted solutions in excess quantity (10 mg), stirred for 72 hrs and filtered through 0.45 μm membrane filter. The filtrate thus obtained was analyzed for AZM content by UV-Visible spectrophotometer [13]. The apparent stability constant (K_1:1_ or K_c_) for the complex was calculated from the slope of the phase-solubility diagrams using the equation;

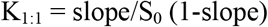

The slope is obtained from the initial straight-line portion of the plot of AZM concentration against SBE-β-CD concentration, and S_0_ is the solubility of AZM in water, in absence of SBE-β CD.

#### 2.2.3 Development and characterization of different formulations

##### 2.2.3.1 Formulation of AZM in situ gel

HPMC (K4M) was added in phosphate buffer (pH 7.4) and was allowed to hydrate. Further, carbopol 934P was sprinkled over this solution and allowed to hydrate over-night (Table 1). Edetate disodium (EDTA) solution was then added to the solution with continuous stirring. Kneaded mixtures of AZM and SB-β-CD were prepared using different molar ratios and added to the carbopol/HPMC solution with continuous stirring until the contents were uniformly blended. Benzalkonium chloride (BKC) was then added and the solution was brought up to 100 ml with addition of purified water. All formulations were allowed to equilibrate for 24 hrs at room temperature prior to further evaluation as shown in Table 1.

**Table 1:** Formulation composition of Azithromycin in situ ophthalmic gel

| Batch code | F1 | F2 | F3 | F4 | F5 | F6 | F7 | F8 | F9 |
| --- | --- | --- | --- | --- | --- | --- | --- | --- | --- |
| AZM: SBE-β-CD ratio | 1:1 | 1:1 | 1:1 | 1:2 | 1:2 | 1:2 | 2:1 | 2:1 | 2:1 |
| AZM (g) | 1 | 1 | 1 | 1 | 1 | 1 | 1 | 1 | 1 |
| Carbopol 934P (g) | 0.4 | 0.6 | 0.8 | 0.4 | 0.6 | 0.8 | 0.4 | 0.6 | 0.8 |
| HPMC K4M (g) | 0.4 | 0.4 | 0.4 | 0.4 | 0.4 | 0.4 | 0.4 | 0.4 | 0.4 |
| Edetate disodium (g) | 0.01 | 0.0<br>1 | 0.0<br>1 | 0.0<br>1 | 0.0<br>1 | 0.0<br>1 | 0.0<br>1 | 0.0<br>1 | 0.0<br>1 |
| Benzalkonium chloride (g) | 0.01 | 0.0<br>1 | 0.0<br>1 | 0.0<br>1 | 0.0<br>1 | 0.0<br>1 | 0.0<br>1 | 0.0<br>1 | 0.0<br>1 |
| Citric acid (g) | 0.40 | 0.4<br>0 | 0.4<br>0 | 0.4<br>0 | 0.4<br>0 | 0.4<br>0 | 0.4<br>0 | 0.4<br>0 | 0.4<br>0 |
| Disodium hydrogen Phosphate (g) | 1.12 | 1.1<br>2 | 1.1<br>2 | 1.1<br>2 | 1.1<br>2 | 1.1<br>2 | 1.1<br>2 | 1.1<br>2 | 1.1<br>2 |
| Purified water (q.s.) | 100 | 10<br>0 | 10<br>0 | 10<br>0 | 10<br>0 | 10<br>0 | 10<br>0 | 10<br>0 | 10<br>0 |

##### 2.2.3.2 Stability study of AZM

Antimicrobial activity of AZM was carried out to check the stability of kneaded AZM:SBE-β-CD inclusion complex. Sterile Nutrient Broth was prepared by using Beef extract (0.3g), peptone (0.3g), sodium chloride (0.1g) and distilled water (30ml). Master culture of *Bacillus subtilis* was prepared in Nutrient Broth and was incubated for 24 hrs at 37°C for the growth of microorganisms.

##### 2.2.3.3 Preparation of AZM solutions

Solution (A) i.e., plain AZM aqueous solution was prepared by adding 0.1 g of AZM in 10 ml of purified water. Solution (B) i.e., AZM aqueous solution containing SBE-β-CD (1:1 molar) was prepared by adding 0.1 g of AZM and 0.28 g of SBE-β-CD in 10 ml of water. Both the solutions were kept in stability chamber at 40°C during the entire period.

##### 2.2.3.4 Preparation of Nutrient Agar plates

Nutrient Agar medium was prepared and transferred aseptically into the two sterilized Petri plates and allowed to solidify. The *B. subtilis* culture was added to the Nutrient Agar plates by pour plate method and incubated at 37°C. Further, the wells were made in the Petri plates in the aseptic cabinet using sterile borer of diameter 10 mm. The solutions (A) and (B) were transferred into the wells and plates were incubated at 37°C. Eventually the zone of inhibition was measured after 2, 7, 15 and 28 days[14].

##### 2.2.3.5 Drug content determination

Accurately 0.5 ml of formulation was pipetted out and diluted with distilled water upto 100 ml. 5 ml aliquot was withdrawn from this solution and diluted to 25 ml distilled water[15]. Finally, AZM concentration was determined at λ_max_= 232 nm using following formula[16],

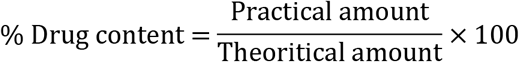

##### 2.2.3.6 Rheological study using Brookfield viscometer

The viscosity of the prepared *in situ* gel was determined using a Brookfield viscometer (DVII Pro digital viscometer) with spindle number 96. The formulation was added to a beaker and was allowed to settle for 30 min at room temperature before the final measurement. Spindle was lowered perpendicularly into the center of *in situ* gel formulation taking care that spindle does not touch bottom of the beaker and rotated at speed of 5, 10, 20, 50 rpm for 5 minutes[17]. The viscosity was recorded at different time points and the average of three readings was recorded with standard deviation.

##### 2.2.3.7 Gelling capacity

In order to determine the gelling capacity 100 μl of the formulation was placed in a vial containing 2 ml of freshly prepared artificial tear fluid (stock solution of NaCl 0.670g, Sodium bicarbonate 0.200g, CaCl_2_.2H_2_O, 0.008g in 100 ml purified water q.s.). The mixture was equilibrated at 37°C and gel formation was assessed visually along with recording the time[18].

##### 2.2.3.8 In vitro drug release study

The *in vitro* release of AZM from the prepared formulation was determined in freshly prepared phosphate buffer through dialysis sac method. The dialysis membrane was first soaked overnight in the dissolution medium; finally 1 ml volume of the formulation was accurately pipette in to this assembly. This membrane was fastened to the metallic drive shaft followed by its suspension in a beaker containing 150 ml of the dissolution medium with temperature maintained at 37±O.5°C. This was done to ensure the continuous contact of dialysis tube with the receptor medium surface. The shaft’s rotation speed was set at 50 rpm; 1 ml aliquots were withdrawn at regular time intervals and replaced by an addition of equal volume to the receptor medium[19]. The aliquots were further diluted with receptor medium and absorbance was measured at 260nm.

##### 2.2.3.9 Stability studies of the optimized in situ gel

The stability studies of prepared ophthalmic *in situ* formulations were carried out at (40°C/75% RH) for one month [20]. Effect of temperature, humidity and time were evaluated on the gel for assessing the stability of the prepared formulation.

##### 2.2.3.10 Eye irritation test

The OECD guideline 405 was followed for *in vivo* eye irritancy test of the developed formulation, which is intended exclusively to be used on albino rabbits. Initially, the test substance was applied in a single dose into the conjunctival sac of one eye of the rabbit. The other eye was left untreated and considered as a control. An *in vivo* eye irritancy confirmatory test was carried out on the basis of the irritant effect in the initial test. According to the guideline, it was recommended to conduct the test in a sequential manner in one animal at a time, rather than exposing two additional animals. Therefore, the irritant or negative response was again confirmed using two additional animals. Finally, the ocular irritation was recorded in terms of scores at 1, 24, 48 and 72 hrs, respectively[21].

## 3. RESULTS AND DISCUSSION

### 3.1 Characterization of inclusion complex of AZM and SBE-β-CD in solid phase using XRD

Powder X-ray diffraction (PXRD) pattern of AZM shows intense sharp peaks at 2Θ at 9.30, 9.811, 18.65,18.75 and 19.13 as shown in Figure 1. The spectrum contains various crystalline peaks for AZM however SBE-β-CD being amorphous in nature shows less intense peaks[22]. A sharp crystalline peak was observed for AZM at 9.30 (Figure 1C) however due to formation of inclusion complex of AZM and SBE-β-CD (Figures 1A and 1B) the peak disappeared as a resultant change in environment. The crystalline peaks of AZM around 16.83, 17.49, and 20.36 were also reduced indicating reduction in its crystallinity. Further, the reduction at 9.811 indicates reduction in crystallinity of SBE-β-CD after inclusion complex formation.

**Figure 1:**
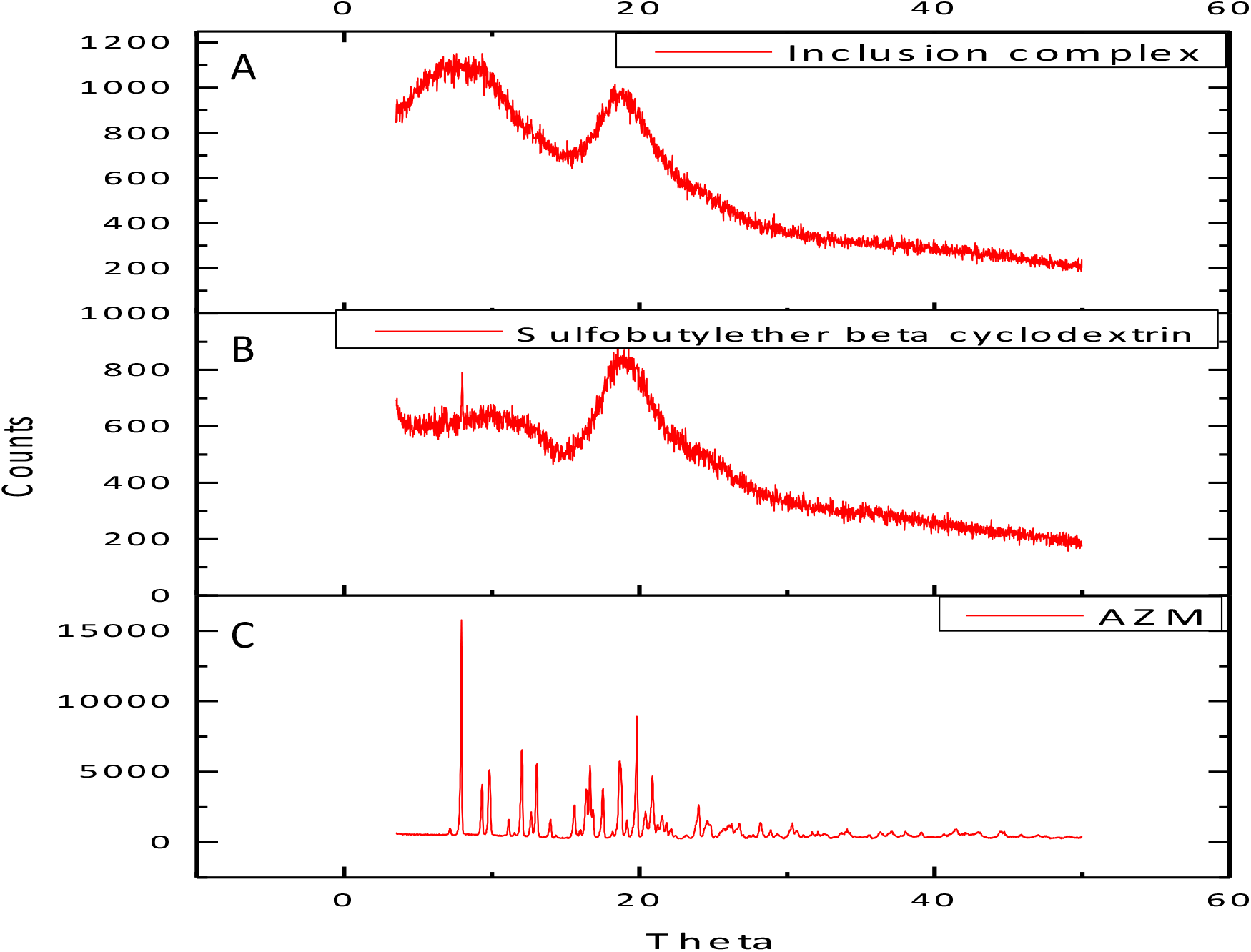
X-ray diffraction pattern of formulations components: (A) AZM:SBE-β-CD inclusion complex (1:1), (B) SBE–β-CD, (C) AZM. Here X-axis corresponds to diffraction angle between incident beam and transmitted beam whereas count on Y-axis represents intensity of the peak.

### 3.2 Inclusion complex characterization study using FTIR spectroscopy

The FTIR spectrum of AZM is characterized by stretching band at 3494.28 cm^-1^for O-H stretching vibrations and 2972.78 cm^-1^ for C-H stretching vibrations (Figure 2). There are strong absorption bands at 1723.40 cm^-1^ for C=O stretching vibration, as well as at the 1187.15 and 1048.07 cm^-1^ bands for C-H and C-O stretching vibrations, respectively[23]. The FTIR spectrum of SBE-β-CD showed a broad of 3421.57 cm^-1^ owing to O-H stretching vibrations. In addition, C-H stretching bands were observed around the region of 2935.82 cm^-1^. A band with a transmittance peak around 1647 cm^-1^ was due to H-O-H bending of water molecules linked with SBE-β-CD while the absorption bands at 1186 and 1043 cm^-1^ was observed for C-H and O=S=O stretching vibrations, respectively[24]

**Figure 2:**
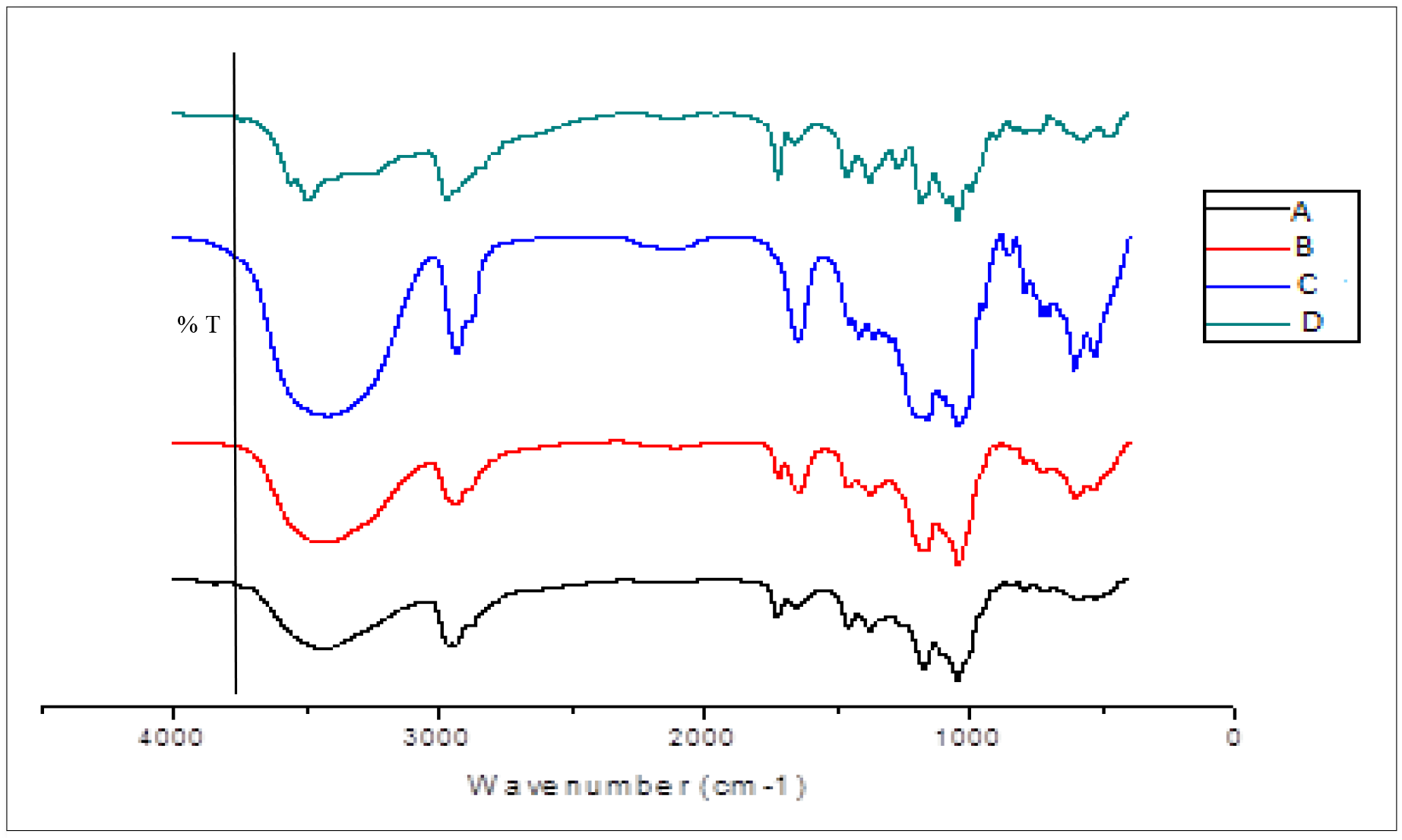
FTIR spectrum of (A) AZM:SBE-β-CD inclusion complex, (B) AZM:SBE-β-CD physical mixture, (C) SBE-β-CD, (D), AZM

The spectra of the AZM:SBE-β-CD inclusion complex show absorption band at 3438.33 cm^-1^for O-H stretching vibration, 2940.69 cm^-1^ for C-H stretching vibration, 1651.80 cm^-1^ for C-N stretching vibration and 1047.77 cm^-1^ for S=O stretching vibration, which implies that stronger interactions existed between AZM and SBE-β-CD. The spectra of the physical mixtures of AZM:SBE-β-CD and AZM:SBE-β-CD inclusion complex are similar in nature and look like a superimposition. The absorption band at 3260 cm^-1^ for O-H stretching vibration from AZM disappeared in the spectrum of the AZM:SBE-β-CD inclusion complex as compared with the spectrum of physical mixture of AZM:SBE-β-CD, which show stronger interactions occurred between AZM and SBE-β-CD (there must be no interaction in physical mixture, not possible at all and if there is interaction, it denotes the formation of cyclodextrin complex).

### 3.3 Dose Calibration of AZM in phosphate buffer (pH 7.4) Using UV visible spectroscopy

The λ_max_ of AZM was found to be 260 nm in phosphate buffer (pH 7.4) as shown in Figure 3A; the slope and intercept were found to be 0.002 and 0.041, respectively with correlation coefficient R^2^ of 0.998. The solubility of AZM in phosphate buffer (pH 7.4) was found to be 0.22 mg/ml. Further, it was observed that 2.27 ml of the dissolution medium was required to dissolve a single dose of AZM (0.5 mg) as shown in Supplementary Table 1.

**Figure 3:**
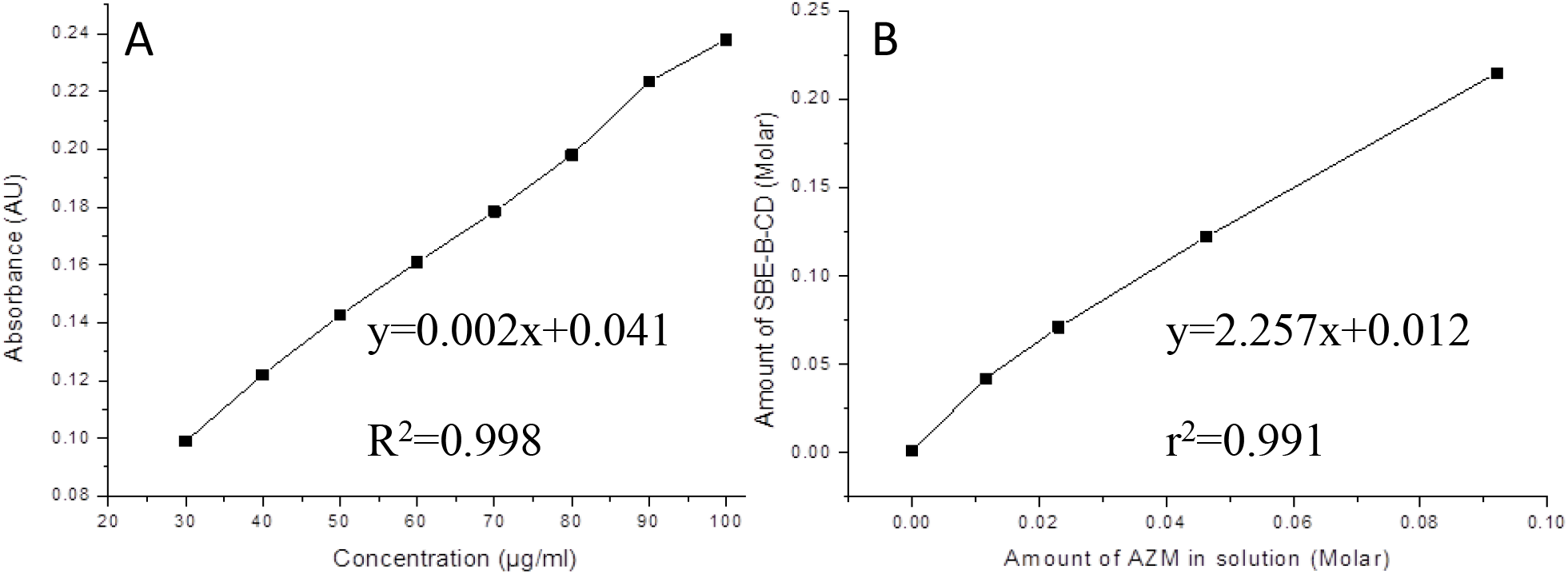
(A) Calibration plot of AZM in phosphate buffer (pH 7.4) was linear in the concentration range of 2 30-100 μg/ml. The statistical R of the plot is 0.998. (B) Phase solubility plot depicts increase in solubility of 2 AZM with increasing concentration of SBE-β-CD having R =0.991.

### 3.3 Solubility enhancement of AZM using Phase solubility study

The phase solubility diagram was obtained at 25 °C by plotting the apparent equilibrium concentrations of the drug against SBE-β-CD concentrations (Figure 3B). The apparent solubility AZM increased linearly as a function of SBE-β-CD concentration (R^2^=0.991) up to 0.1 M, corresponding to the aqueous solubility of SBE-β-CD. The linear relation between AZM solubility and SBE-β-CD concentration indicates a AL-type phase-diagram, defined by Saokham *et al.* [25]. This diagram is characteristic of 1:1 complexation and suggested that water-soluble complex was formed in solution. The slope of the curve (m) was found to be 2.257 whereas the intercept (c) was 0.012 along with the correlation coefficient of 0.991 as shown in Figure 3B. The phase solubility studies showed that SBE-β-CD increases solubility of drug linearly. Phase solubility plot is almost linear and calculated value of stability constant K_1:1_ was found to be 90.45 M^-1^, depicting the strength of interaction between AZM and SBE-β-CD, and eventually confirming the formation of the inclusion complex.

### 3.4 Stability study of AZM in presence of SBE-β-CD and its antimicrobial characterization

The physical mixture showed similar inhibitory activity as it was observed for free AZM, whilst the isolated SBE-β-CD showed no antibacterial effect. Further, the microbiological study depicted that the AZM:SBE-β-CD inclusion complexes had the same inhibitory effect as the free AZM, indicating that the drug stability was maintained throughout the preparation methods in this study. Antimicrobial study indicates no significant difference in zone of inhibition of plain AZM aqueous solution and its aqueous solution with SBE-β-CD (Figure 4A). Moreover, zones of inhibition of AZM aqueous solution decreased significantly with time (t-test, p<0.05), whereas the zones of inhibition of AZM aqueous solution with SBE-β-CD remained unchanged. This indicates the stabilizing effect of SBE-β-CD on AZM as shown in Figure 4B. Thus, from the above study it was observed that inclusion of AZM in cyclodextrin improved the storage stability and also had no inhibitory effect on antimicrobial activity when compared to AZM pure drug.

**Figure 4:**
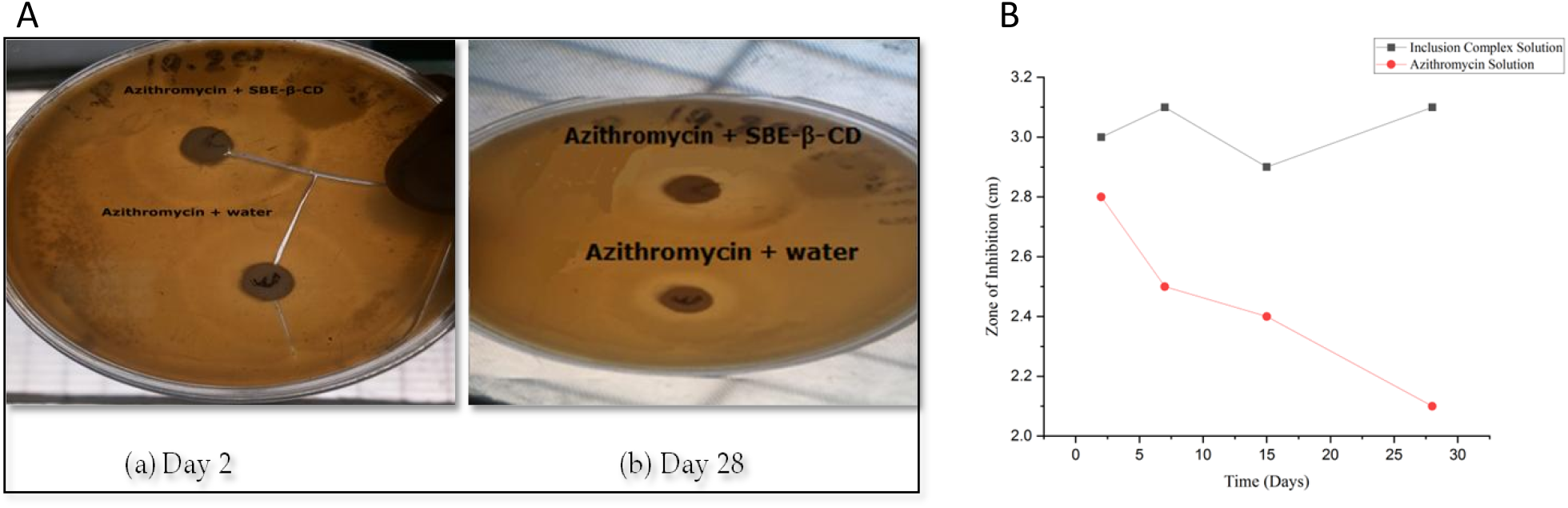
(A) Photographic image of nutrient agar plate portrays the zone of inhibition (a) Azithromycin solution and Physical mixture of AZM and SBE-β-CD, (b) AZM:SBE-β-CD inclusion complex and Azithromycin solution after day 2 and day 28. (B) Graphical representation for zone of inhibition v/s time of AZM:SBE-β-CD inclusion complex (black color filled square dots) and AZM solution (red color filled circles)

### 3.5 Measurement of gelling capacity and determination of drug content of AZM *in situ* gel

The AZM content of all formulations was found to be ranging between 85.9±8.192 to 95.2±4.25 (Table 2). The drug content of all formulation was found within the compendia specification limit of 85%-115%. This is outstanding observation for these formulations sees in *in situ* gel. The gelling capacity data of prepared formulations presented in Table 2 show that the formulation F2 underwent prompt gelation and rapid dissolution. On the contrary the formulation F8 had immediate gelation but existed only for one hour. This connoted that the transition time of formed gel was reduced whereas the erosion time was increased on increasing the polymer concentration.

**Table 2:** Gelling capacity and % drug content of prepared formulations of AZM in situ gel

| Formulation Code | Gelling capacity | Result | % Drug content (Practical content/Theoretical) *100 |
| --- | --- | --- | --- |
| F1 | - | No gelation. | 95.2±4.25 |
| F2 | + | Gel formation after a few minutes, dissolves rapidly | 89.7±2.532 |
| F3 | ++ | Instant gelation, remains for few hours | 92.2±3.619 |
| F4 | +++ | Immediate gel formation and stable for longer period | 93.1±3.05 |
| F5 | ++ | Instant gelation, remains for few hours | 94.3±3.883 |
| F6 | ++ | Instant gelation, remains for few hours | 93.6±6.686 |
| F7 | ++ | Instant gelation, remains for few hours | 85.9±8.192 |
| F8 | +++ | Immediate gel formation and stable for longer period | 94.2±3.758 |
| F9 | + | Gel after a few minutes, dissolves rapidly | 91.7±2.91 |

### 3.6 Rheological study of AZM *in situ* gel

Newtonian flow was exhibited by all the formulations on rheological evaluation before gelling while pseudoplastic flow was seen for these formulations after gelling in the eye. There was increase in the viscosity after gelling as seen in our case. Additionally, the gel formed *in situ* retained its integrity for an extended period of time without dissolving or eroding. The results of rheological studies of *in situ* gel showed that the viscosity decreased as the shear stress (rpm) increased. With this increase in shear stress, the normally disarranged molecules of the gelling material aligned their long axis in the direction of the flow (Figure 5). Such orientation reduced the internal resistance of the material and hence, decreased the viscosity. Figure 6A shows the average effect of AZM:SBE-β-CD ratio on viscosity of the formulation. This graph shows that the viscosity of *in situ* gel decreases with the increasing percentage of SBE-β-CD in respect to the drug (1:1). Further increase in percentage of SBE-β-CD elevates the viscosity of the formulation.

**Figure 5:**
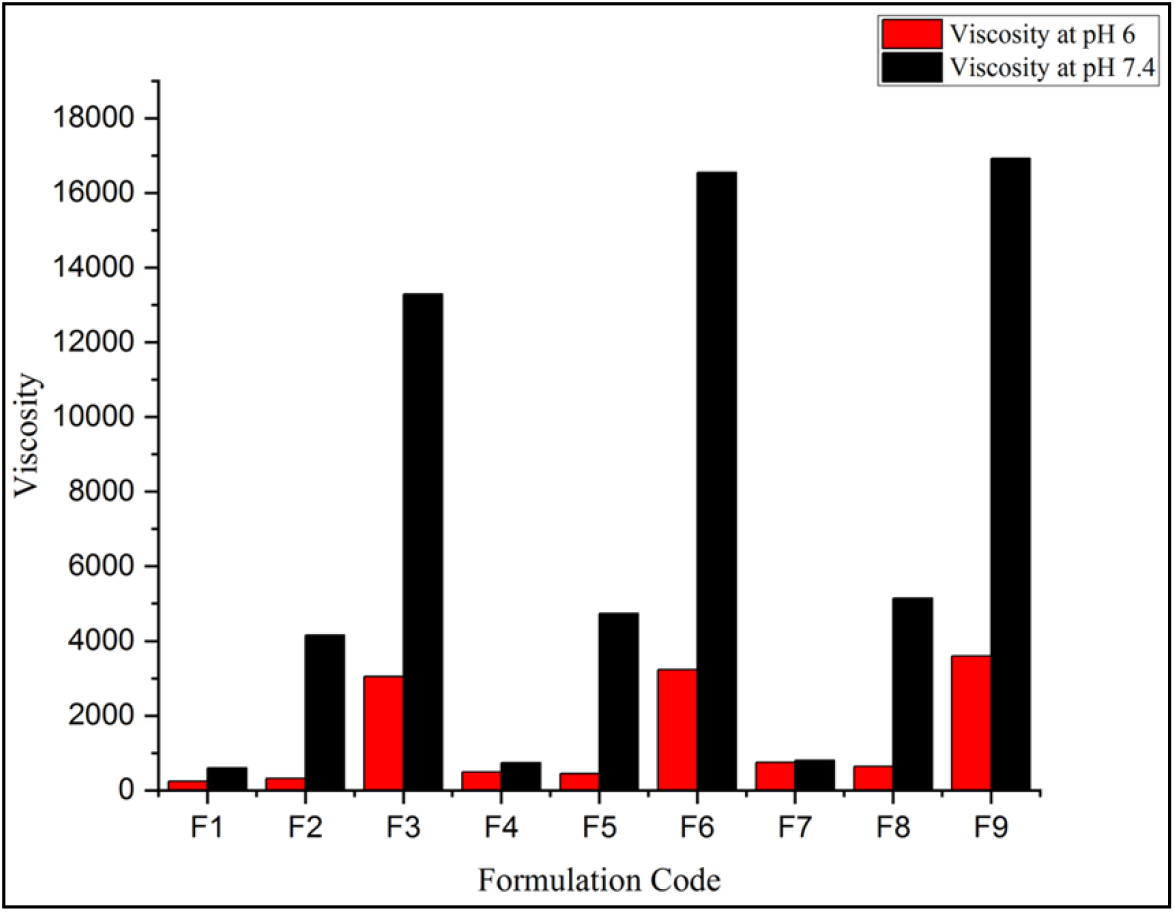
Graphical representation of viscosity of prepared formulations at pH 6.0 (solution phase in red color bars) and pH 7.4 (gel phase in black color bars) of in situ gel

The average effect of concentration of carbopol 934P on viscosity of the formulation is shown in Figure 6B. At pH 6, there was practically no significant effect of the concentration of carbopol 934P (up to 0.6%) but further increase in its concentration leads to the high increase in viscosity of the formulation. At pH 7.4, viscosity was increased with increase in amount of carbopol 934P (from 0.4g to 0.8 g). The addition of carbopol led to formation of an *in situ* viscous gel that minimized the leakage of solution from the eye during instillation[26]. Thus, it was observed that at pH 7.4 (physiological pH of eye) all the formulation transitioned to gel form thereby confirming the pseudoplastic nature of *in situ* gel. Further, Gelling capacity of these formulations is mentioned in Table 2 and Figure 6C.

**Figure 6:**
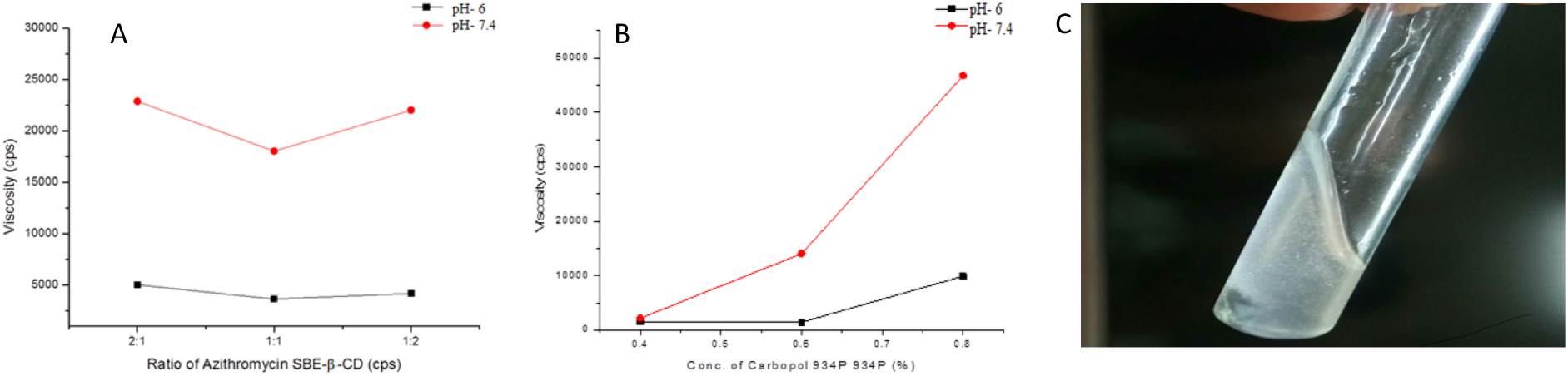
(A) Graph showing the average effect of concentration of AZM:SBE-β-CD ratio on viscosity at 20 rpm at pH 6.0 (solution phase; black color) and pH 7.4 (gel phase; red color) (B) Graph showing the average effect of concentration of Carbopol 934P on viscosity at 20 rpm at pH 6.0 (solution phase; black color) and pH 7.4 (gel phase; red color) (C) Formation of AZM ophthalmic in-situ gel

### 3.7 *In vitro* drug dissolution study of AZM *in situ* gel

The *in vitro* drug release of AZM *in situ* gel was carried out in the dissolution testing apparatus. Formulation (F4) showed the highest drug release of 79.5±6%as shown in Supplementary Table 2 and Figure 7. The highest drug release in formulation F4 may be due to the higher amount of SBE-β-CD and lower amount of carbopol 934P. Lower amount of carbopol934P led to the formation of less viscous gel and showed better drug release.

**Figure 7:**
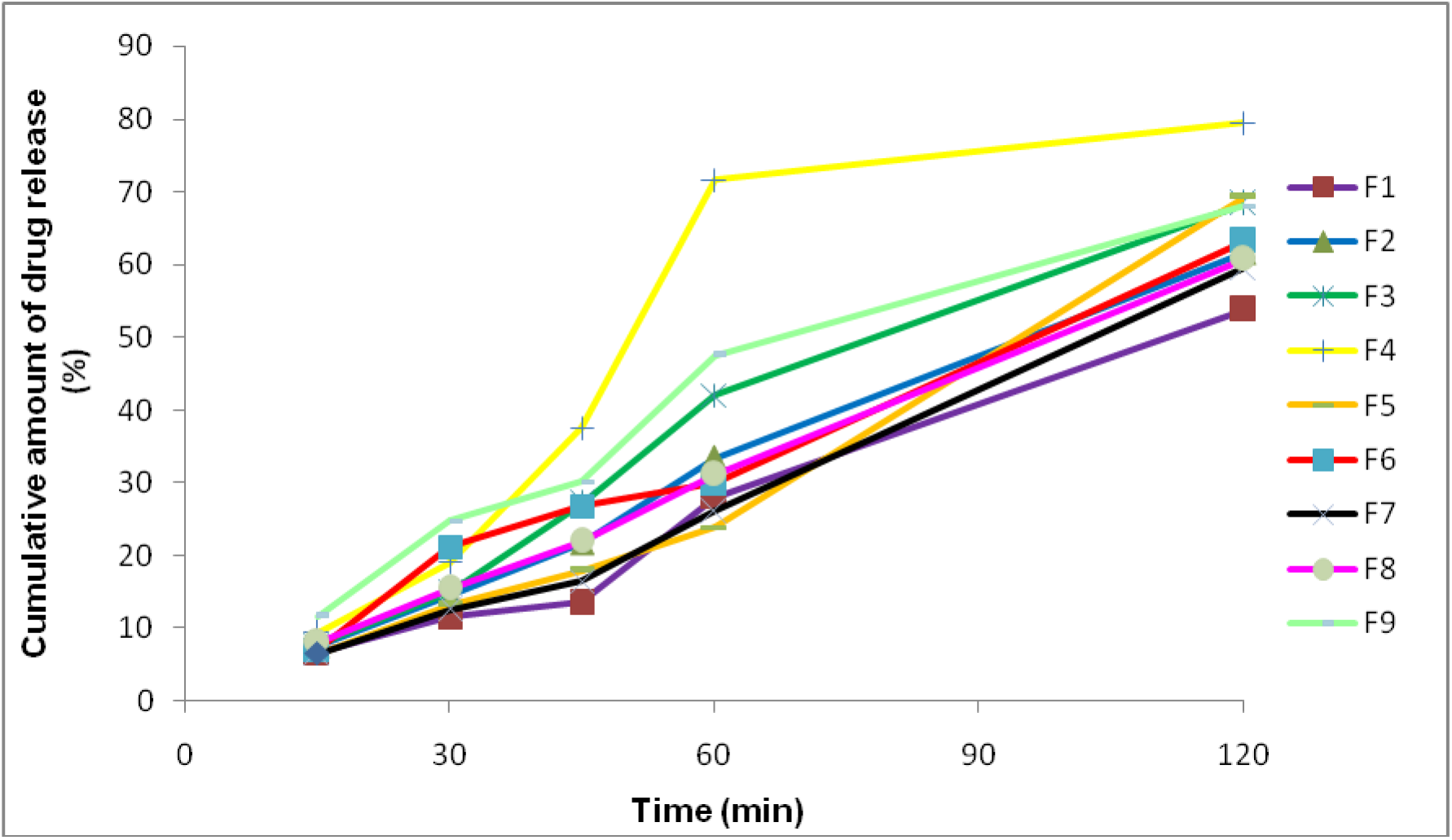
Graph portrays cumulative in-vitro release profile of prepared AZM in situ gel formulations in phosphate buffer (pH 7.4) at 37°C

### 3.8 *In vitro* stability study of the optimized *in situ* gel

The optimized sterilized formulation showed good physical stability and no discoloration or physical changes were observed after storage as shown in Table 3; Supplementary Table 3 and Supplementary Figure 1.

**Table 3:** Stability parameters of the, formulation F4 in situ ophthalmic gel of AZM

| Quality attribute | Sampling intervals |  |  |  |  |
| --- | --- | --- | --- | --- | --- |
|  | 0 week | 1<sup>st</sup> week | 2<sup>nd</sup> week | 3<sup>rd</sup> week | 4<sup>th</sup> week |
| pH | 6.3 | 6.3 | 6.3 | 6.3 | 6.3 |
| Clarity | Clear | Clear | Clear | Clear | Clear |
| Drug content | 92.1±3.05 | 91.4±3.02 | 90.4±3.02 | 89.8±2.04 | 88.9±2.03 |
| Gelling capacity | +++ | +++ | +++ | +++ | +++ |

### 3.9 Ocular irritancy test of AZM *in situ gel*

The eye irritation studies were carried out to evaluate the tolerability of the *in situ* gel. The eye study conducted on rabbit by instilling of gel for 3 days did not show any sign of irritation i.e., 0 score for all grades, implying that rabbits tolerated the optimized checkpoint formulation very well (Table 4 and Figure 8).

**Figure 8:**
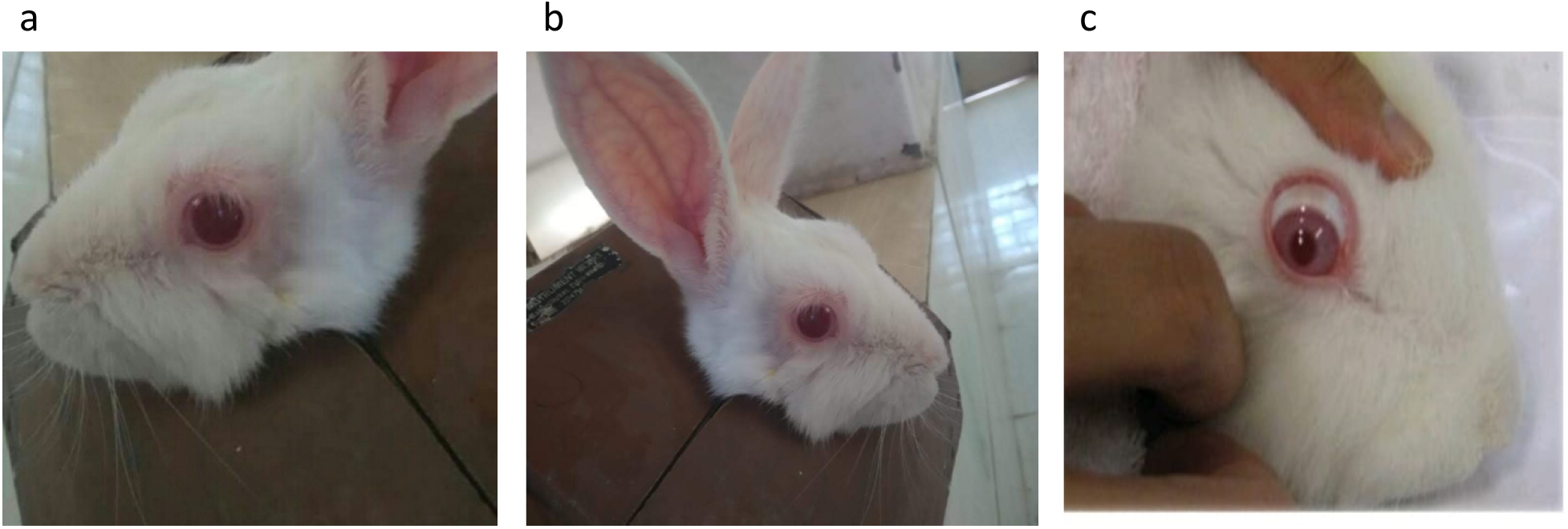
Photographs of rabbit used for ocular irritation test (a) for Control group, (b) after 1 hour and (b) after 72 hours with no sign of redness and irritation

**Table 4:** The grading response for the ocular irritation test on rabbit eye at different time intervals

| Time (hrs) | Redness of eye | Itching of eye | Uveitis | Epiphora |
| --- | --- | --- | --- | --- |
| 1 | 0 | 0 | 0 | 0 |
| 4 | 0 | 0 | 0 | 0 |
| 24 | 0 | 0 | 0 | 0 |
| 48 | 0 | 0 | 0 | 0 |
| 72 | 0 | 0 | 0 | 0 |

## 4. CONCLUSION

The poorly soluble drug AZM was successfully formulated as *in situ* gelling ophthalmic formulation using carbopol 934P as a pH-triggered gelling agent. The solubility and stability of AZM was improved using SBE-β-CD. The phase solubility study clearly indicates increase in solubility of AZM on addition of SBE-β-CD. Antimicrobial study indicated the reduction in degradation of AZM on addition of SBE-β-CD. The gelling and viscosity studies indicated the solution to gel transition and subsequent increase in viscosity of the formulation on changing the pH of the medium. Further, *in vitro* release studies showed the extended release of AZM within a period of 2 hours. Considering the gelling and release studies, formulation F4 was selected as best candidate for further studies. The stability study and eye irritation study showed the stability and compatibility of the formulation with ocular tissues. Thus, the developed *in situ* gelling ophthalmic formulation of AZM can be a viable alternative to conventional eye drops of AZM which suffer from demerits of poor solubility, poor stability, rapid pre-corneal elimination of the formulation and a need for high frequency of eye drop instillation. These results will open up further discussion for pharmacological intervention with SBE-β-CD and provide a potentially strong alternative for treatment of ophthalmic infections.

## Abbreviations

AZM: Azithromycin
SBE-β-CD;: sulfobutylether β-cyclodextrin
HPMC: hydroxypropyl methylcellulose

## Funding Source

No funding declared

## Conflict of Interest

Authors declare no conflict of interest.

## Acknowledgements

Authors are thankful to Sun Pharma Paonta Sahib, and Lachoo Memorial College of Science and Technology (Autonomous), Jodhpur for their support throughout the research work.

